# Performance of the cobas^®^ HBV RNA Automated Investigational Assay for the Detection and Quantification of Circulating HBV RNA in Chronic HBV Patients

**DOI:** 10.1101/2022.03.09.483670

**Authors:** Caroline Scholtès, Aaron T. Hamilton, Marie-Laure Plissonnier, Caroline Charre, Beth Scott, Ling Wang, Françoise Berby, Janine French, Barbara Testoni, Alan Blair, Miroslava Subic, Matthias Hoppler, Andreas Lankenau, Andreas Grubenmann, Massimo Levrero, Marintha L. Heil, Fabien Zoulim

**Author notes:** Corresponding author, Cancer Research Center of Lyon, UMR Inserm 1052 - CNRS 5286, 151 cours Albert Thomas, 69424 Lyon Cedex 03.

## Abstract

**Background:** The amount of HBV RNA in peripheral blood may reflect HBV covalently closed circular DNA (cccDNA) transcriptional activity within infected hepatocytes. Quantification of circulating HBV RNA (cirB-RNA) is thus a promising biomarker for monitoring antiviral treatment.

**Objectives:** We evaluated the performance of an automated, prototype quantitative HBV RNA assay for use on the Roche **cobas**^®^ 6800/8800 systems.

**Study Design:** The sensitivity, specificity, linearity, and potential interference by HBV DNA of the **cobas**^®^ HBV RNA assay were assessed using synthetic HBV armored RNA and clinical specimens.

**Results:** **cobas**^®^ HBV RNA results were linear between 10 and 10^7^ copies/mL in clinical samples of several HBV genotypes, and up to 10^9^ copies/mL with synthetic RNA. Precision and reproducibility were excellent, with standard deviation below 0.15 log_10_ copies/mL and coefficients of variation below 5% throughout the linear range. The presence of HBV DNA had minimal (<0.3 log_10_ copies/mL) impact on HBV RNA quantification at DNA:RNA ratios of up to approximately one million. In a panel of 36 untreated patient samples, cirB-RNA concentrations were approximately 200-fold lower than HBV DNA. cirB-RNA was detected in all 13 HBeAg-positive patients (mean 6.0 log_10_ copies/mL), and in 20 of 23 HBeAg-negative patients (mean of quantifiable samples 2.2 log_10_ copies/mL). Finally, cirB-RNA was detected in 12 of 20 nucleoside analog-treated patients (mean of quantifiable samples 3.4 log_10_ copies/mL).

**Conclusions:** The **cobas**^®^ 6800/8800 investigational HBV RNA assay is a high throughput, sensitive and inclusive assay to evaluate the clinical relevance of cirB-RNA quantification in patients with chronic hepatitis B.

## Background

Hepatitis B virus (HBV) circulating RNA (cirB-RNA) is a promising biomarker for definition of antiviral treatment endpoints, since circulating pregenomic RNA (pgRNA) has been proposed to reflect the pool of transcriptionally active covalently closed circular DNA (cccDNA) within infected hepatocytes. Previous publications have indicated that serum HBV RNA levels have good predictive power for both on-treatment serologic response and off-treatment durability [1–4]. Moreover, the combination of undetectable cirB-RNA and HBV core-related antigens (HBcrAg) at the end of treatment was shown to have a better predictive value for off-treatment outcomes than either biomarker alone [1]. In the context of emerging antiviral agents [5], robust assays with high sensitivity and accuracy over a broad linear range are crucial for assessment of antiviral drug mechanisms of action, the impact of the drug on cccDNA transcriptional activity, and the ability to predict the achievement of treatment endpoints [6, 7]. However, no standardized assay for quantifying cirB-RNA exists, which hampers widespread application of cirB-RNA quantification in the clinical management of chronic hepatitis B (CHB) patients. The majority of currently available tests have a lower limit of quantification (LLOQ) around 10^3^ copies/mL, although in-house RT-droplet digital PCR (ddPCR) assays [8] and the Abbott serum HBV pgRNA assay [9] have LLOQ of approximately 10^2^ copies/mL. This limitation might compromise the diagnostic performance of this method, particularly among HBeAg-negative patients who often present with very low cirB-RNA levels.

## Objectives

In this study we assessed the analytical and clinical performance of the investigational **cobas**^®^ HBV RNA assay (cobas HBV RNA) on the **cobas**^®^ 6800/8800 System.

## Study Design

### Synthetic RNA and DNA templates

Synthetic armored RNA (arRNA) containing 435 bp derived from the 3′ end of HBV pgRNA, packaged in MS2-phage [10] was used for assay performance evaluation. arRNA was quantified by ddPCR (BioRad) using primers and a probe in the precore/core region. Synthetic HBV DNA, encompassing the same HBV sequence as the arRNA, was packaged in a lambda phage vector [11]. HBV DNA was quantified using **cobas**^®^ HBV for use on the **cobas**^®^ 6800/8800 Systems (Roche Molecular Diagnostics, Pleasanton, CA) which has an LLOQ of 10 international units (IU)/mL.

### Patient samples

Clinical samples were from 56 patients included in the ANR-17-RHUS-0003 cirB-RNA cohort [12], who provided written informed consent (see Supplemental Material for details of ethics considerations and Table S1 for patient characteristics). Twenty of these patients were treated with nucleoside analogs (NUC) tenofovir or entecavir. A subset of the 56 patient samples was used for linearity (Table S2) and method comparison experiments. HBV genotypes were determined using the ViroKey SQ FLEX Genotyping Assay (Vela Diagnostics, Hamburg, Germany). The samples were tested for HBsAg, HBeAg (Abbott Diagnostics, Des Plaines, IL, USA) and HBV DNA (**cobas**^®^ HBV).

### cirB-RNA measurement

The Roche HBV RNA investigational assay for use on the **cobas**^®^ 6800/8800 Systems (cobas HBV RNA, Roche Diagnostics, Pleasanton, CA) quantifies HBV RNA in EDTA plasma or serum. The assay includes an internal control for nucleic acid recovery. The amplification target is located at the 3′ end of HBV transcripts (Figure 1A), enabling it to detect all viral RNAs expressed from cccDNA. *In-silico* sequence analysis indicates that the assay is expected to perform equivalently on all HBV genotypes. Up to 93 samples can be tested in 3.5 hours on the **cobas**^®^ 6800/8800 System. All tests in the present study were performed using the **cobas**^®^ 6800 by trained operators according to the manufacturers’ specifications. The cobas HBV RNA assay is not approved for clinical use by any regulatory body.

**Figure 1.**
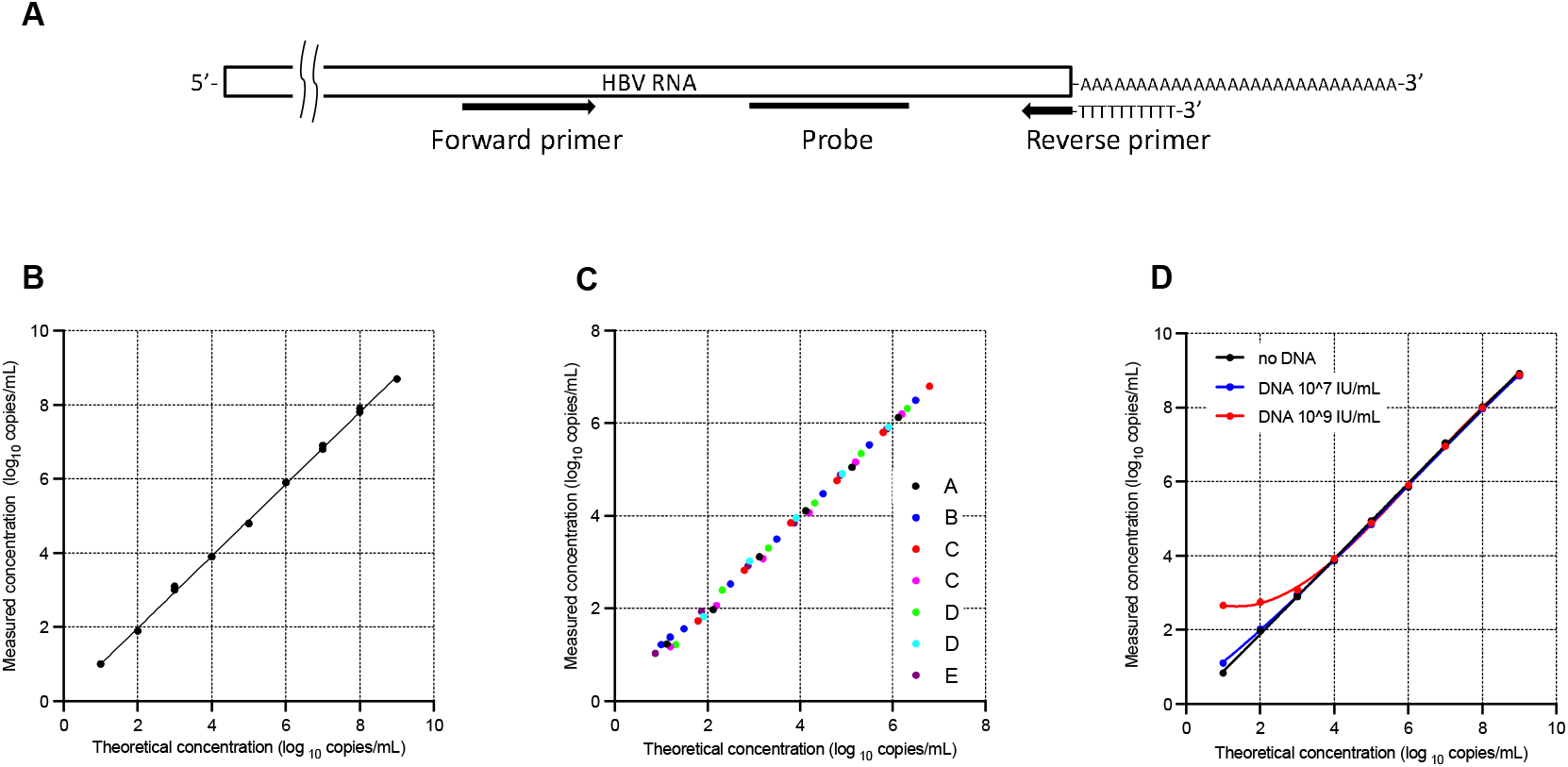
cobas HBV RNA assay design and performance. A. Schematic diagram of primers and probe used in cobas HBV RNA. B. Linearity assessed using 10-fold serial dilutions of arRNA from 10 to 10^9^ copies/mL. C. Linearity assessed using patient samples representing HBV genotypes A, B, C (2 patients), D (2 patients), and E diluted in negative plasma. Slopes of regression lines ranged from 0.97 (genotype E) to 1.01 (genotype C, patient 1), Y-intercepts ranged from −0.12 (genotype C, patient 1) to 0.17 (genotype B), and R^2^ values were all higher than 0.998. D: Impact of high HBV DNA concentration on HBV RNA quantification. HBV arRNA was diluted in series from 10^9^ to 10 copies/mL in EDTA-plasma without (black) or with synthetic HBV DNA at fixed concentrations (red: 10^9^ IU/mL, blue: 10^7^ IU/mL).

cobas HBV RNA is calibrated in units of copies/mL, based on an arRNA Roche internal standard quantitated by ddPCR. One copy of RNA is defined to represent a similar number of RNA molecules as an international unit for DNA molecules, with awareness of an inherent uncertainty. This one-to-one equivalence between DNA IU/mL and RNA copies/mL is helpful for comparisons between DNA and RNA concentrations.

### Analytical sensitivity

The limit of detection (LOD) of cobas HBV RNA was assessed using arRNA diluted in EDTA-plasma or serum to concentrations ranging from 1.25 to 20 copies/mL (forty-two replicates per concentration).

### Linearity

arRNA was diluted in EDTA-plasma to target concentrations between 10 and 10^9^ copies/mL and tested in duplicate. Linearity was also assessed using serial dilutions of seven patient samples with high cirB-RNA loads including HBV genotypes A, B and E (one patient each), as well as C and D (two patients each, with low and high DNA concentration).

### Precision and reproducibility

Precision was evaluated using three dilutions of arRNA in plasma at approximately 10^2^, 10^4^, and 10^7^ copies/mL, with 15 repeats each in the same run. Reproducibility was evaluated using two dilutions of the arRNA, at approximately 10^3^ and 10^6^ copies/mL, on 20 different days.

### Analytical Specificity

Specificity was assessed using 20 HBV-negative serum and plasma samples and 28 remnant samples containing human immunodeficiency virus type 1 (n=13), hepatitis C virus (n=10), or hepatitis E virus (n=5) (see Supplemental Material for details). Interference by HBV DNA was evaluated by spiking negative plasma with 50 copies/mL of arRNA (5 X LLOQ) and synthetic HBV DNA at concentrations from 10^3^ to 10^7^ IU/mL.

### Method comparison

HBV RNA concentrations measured by cobas HBV RNA were compared to those from an in-house ddPCR assay using primers and probes targeting the 3′ end of HBV RNA transcripts (see Supplemental Material for details).

## Results

### Analytical sensitivity

The LOD based on arRNA was estimated to be 3.3 copies/mL (95% confidence interval: 2.6 to 4.8 copies/mL) using PROBIT analysis (95% reactive rate) and 5.0 copies/mL by hit rate. Results were equivalent for plasma and serum (Table 1). The LOD was confirmed in clinical samples representing HBV genotypes A, B, C, D, E (Table S3).

**Table 1.** Analytical sensitivity.

| HBV RNA concentration (copies/mL) | N positive/N valid replicates (plasma) | % positive (plasma) | N positive/N valid replicates (serum) | % positive (serum) |
| --- | --- | --- | --- | --- |
| 20 | 84/84 | 100 | 84/84 | 100 |
| 15 | 84/84 | 100 | 84/84 | 100 |
| 10 | 83/83 | 100 | 84/84 | 100 |
| 5 | 82/84 | 97.6 | 83/84 | 98.8 |
| 2.5 | 75/83 | 90.4 | 75/84 | 89.3 |
| 1.25 | 65/84 | 77.4 | 49/84 | 58.3 |
| 0 | 0/84 | 0 | 0/84 | 0 |
| LOD by PROBIT analysis (95% Reactive Rate) | 3.3 copies/mL (95% CI: 2.6 – 4.8 copies/mL) |  | 3.3 copies/mL (95% CI: 2.7 – 4.5 copies/mL) |  |
| LOD by Hit Rate | 5 copies/mL (97.6%) |  | 5 copies/mL (98.8%) |  |

### Linearity

The dynamic range of quantification using arRNA was 10 to 10^9^ copies/mL (slope 0.98, Y-intercept - 0.036, R^2^ 0.9992; Figure 1B). Results from clinical samples were linear between 10 and 10^7^ RNA copies/mL for all genotypes tested (Figure 1C). These data also establish the lower limit of quantification (LLOQ) as 10 copies/mL, the lower end of the linear range.

### Precision and reproducibility

Assay precision was high, with a coefficient of variation of 4.7% for the low concentration sample (~2 log_10_ copies/mL) and <0.7% for higher concentrations (Table 2, Figure S1). All replicates gave a result that was less than 0.2 log_10_ copies/mL different from the median, and ≥95% of results were within 0.08 to 0.27 log_10_ copies/mL of each other. For reproducibility the coefficient of variation was 3.1% at 3 log_10_ copies/mL and 2.3% at 6 log_10_ copies/mL, with standard deviation below 0.15 log_10_ copies/mL (Table 2, Figure S1). Only one replicate (out of 21 replicates at 6 log_10_ copies/mL) gave a result that was more than 0.2 log_10_ copies/mL different from the median, and ≥95% of results were within 0.36 log_10_ copies/mL of each other.

**Table 2.** Precision and reproducibility.

|  | Target RNA level<br>(log<sub>10</sub> copies/mL) | N<br>replicates | Measured<br>concentration (SD)<br>(mean log<sub>10</sub><br>copies/mL) | Coefficient of<br>variation (%) | Range covering<br>95% of results<br><sup>a</sup>(log<sub>10</sub> copies/mL) |
| --- | --- | --- | --- | --- | --- |
| Precision | 2.0 | 15 | 1.91 (0.09) | 4.7 % | 0.27 |
|  | 4.0 | 15 | 3.89 (0.03) | 0.7 % | 0.08 |
|  | 7.0 | 15 | 6.86 (0.04) | 0.6 % | 0.15 |
| Reproducibility | 3.0 | 21 | 2.94 (0.09) | 3.1 % | 0.27 |
|  | 6.0 | 20 | 5.71 (0.14) | 2.4 % | 0.36 |

<sup>a</sup> difference between 2.5 – 97.5 percentiles

### Analytical Specificity

cirB-RNA was undetectable in samples lacking HBV. HBV RNA concentrations measured in plasma and serum samples from HBV infected patients were equivalent (Figure S2).

Specificity for HBV RNA was assessed by adding HBV DNA at high concentrations (from 10^3^ to 10^7^ IU/mL) to samples with cirB-RNA concentrations around the LLOQ (50 copies/mL). Measured HBV RNA concentrations were higher by only 0.03 to 0.09 log_10_ copies/mL in the presence of added DNA (Table 3).

**Table 3.** Impact of varying concentrations of HBV DNA on low concentration HBV RNA quantification.

| HBV arRNA concentration ($\log_{10}$ copies/mL) | HBV DNA concentration ($\log_{10}$ IU/mL) | Observed RNA concentration<sup>a</sup> ($\log_{10}$ copies/mL) | Difference (mean observed - no DNA reference) |
| --- | --- | --- | --- |
| 1.70 | 0 | 1.81 | n/a |
| 1.70 | 3.0 | 1.85 | 0.04 |
| 1.70 | 4.0 | 1.87 | 0.06 |
| 1.70 | 5.0 | 1.84 | 0.03 |
| 1.70 | 6.0 | 1.89 | 0.08 |
| 1.70 | 7.0 | 1.90 | 0.09 |

<sup>a</sup> mean concentration from 15 replicates

We also assessed the impact of adding exogenous HBV DNA to dilutions of arRNA. With 7.0 log_10_ IU/mL HBV DNA, there was no effect on cirB-RNA quantification (observed – expected RNA concentration <0.1 log_10_ copies/mL) when the RNA concentration was 2 log_10_ copies/mL or higher, and a minimal effect (0.27 log_10_ copies/mL) at 1.0 log_10_ copies/mL, where the DNA to RNA ratio was 10^6^ (Figure 1D). With HBV DNA at 9.0 log_10_ IU/mL, cobas HBV RNA was unaffected at 4 log_10_ copies/mL or higher, and affected slightly (0.19 log_10_ copies/mL) at 3 log_10_ copies/mL where the DNA to RNA ratio was 10^6^ (Figure 1D). At the highest DNA:RNA ratio (10^8^), the difference was 1.83 log_10_ copies/mL).

Finally, we assessed the extent to which HBV DNA interferes with cirB-RNA quantification in the serially diluted clinical samples previously described (Figure 1C). Measured and expected cirB-RNA concentrations were not statistically significantly different (mean difference 0.004 log_10_ copies/mL, P value 0.73) regardless of dilution factor and resulting HBV DNA concentration. Notably, in the two samples from NUC-treated patients with relatively low DNA concentrations (one each for genotype C and D; see Table S2), the difference between the measured and expected RNA concentration at dilutions where the expected DNA concentration was below 10 IU/mL (0.005 log_10_ copies/mL) was essentially the same as in dilutions with expected DNA concentrations higher than 10 IU/mL (−0.009 log_10_ copies/mL; P value 0.88) (data not shown).

### Method comparison

arRNA concentrations measured with cobas HBV RNA and an in-house ddPCR assay were highly correlated (Figure S3).

For research purposes that do not require high throughput and automation, a manual version of this assay was developed. Results obtained with this manual assay were highly correlated with those obtained with cobas HBV RNA (Figure S4).

### Evaluation of patient samples

We measured cirB-RNA levels in 56 clinical samples, from 36 untreated and 20 NUC-treated, HBV-infected patients. The samples were selected for their genotype representativity (A to G) and to cover a wide HBV DNA concentration range within the different phases of HBV disease [13]. HBV DNA levels ranged from undetectable to 8.8 log_10_ IU/mL (mean of the quantifiable samples 4.9 log_10_ IU/mL; Figure S4).

Among the 36 untreated patients, all HBeAg(+) patients were cirB-RNA positive with quantifiable values (Figure 2). Mean cirB-RNA levels in untreated HBeAg(+) CH and CI patients were 5.6 log_10_ copies/mL and 6.2 log_10_ copies/mL, respectively (P value 0.21, t-test). cirB-RNA could be quantified in 11 of 17 (65%) untreated HBeAg(-) CH and one of six (17%) HBeAg(-) CI patients. In samples with quantifiable cirB-RNA, mean cirB-RNA levels were higher in HBeAg(+) vs HBeAg(-) patients (6.0 log_10_ vs 2.2 log_10_ copies/mL; p<0.0001). Amongst 33 samples with both RNA and DNA levels over the LLOQ, mean cirB-RNA concentrations were 2.4 log_10_ copies/mL lower than mean HBV DNA levels, with no significant difference between disease phases or genotypes. The difference between DNA and RNA concentrations in untreated patients ranged from 1.5 to 3.4 log_10_ copies/mL.

**Figure 2.**
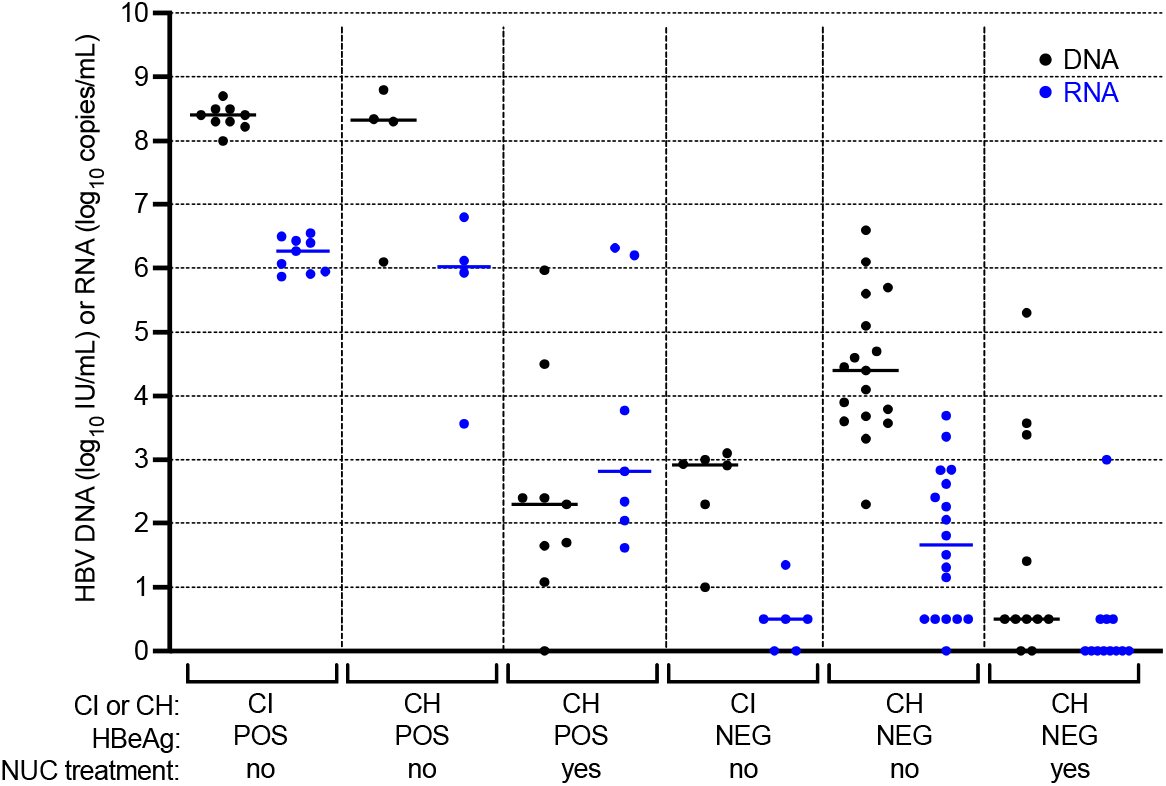
Quantification of HBV cirB-RNA and DNA in clinical samples. Plasma HBV DNA viral load (black) and HBV cirB-RNA (blues) levels in 56 patients plotted according to EASL disease phase, HBeAg seropositivity, and NUC treatment status. For illustrative purposes HBV DNA and cirB-RNA values < LOD were assigned a value of 0 IU or copies/mL, and those < LLOQ but > LOD were assigned a value of 0.5 log_10_ IU or copies/mL. CI: chronic infection; CH: chronic hepatitis. See Supplemental Material Figure S5 for representation of the different HBV genotypes included.

In the 20 NUC-treated patients, the mean of 12 DNA concentrations above the LLOQ was 3.0 log_10_ IU/mL. cirB-RNA was detected in 12 patients; the mean RNA concentration of the nine results above the LLOQ was 3.4 log_10_ copies/mL. Two of these 12 patients had undetectable DNA viral load and five had higher cirB-RNA than HBV DNA concentration.

## Discussion

Several reports have highlighted the potential of cirB-RNA quantification to serve as a surrogate marker for intrahepatic cccDNA transcriptional activity [14, 15] and assessment of antiviral efficacy [16–18]. In a multicenter prospective cohort study, non-cirrhotic patients with undetectable HBV DNA and cirB-RNA at the end of NUC treatment had significantly lower risk of viral relapse in long-term follow-up compared with those who had detectable serum HBV DNA or RNA [2]. Therefore, a sensitive and reliable method for detection of HBV RNA may assist decisions about when to stop NUC therapy.

In recent years, various RT-qPCR-based quantitative methods for serum HBV RNA have been developed, usually with a LLOQ around 1000 copies/mL. The use of digital PCR allowed the reduction of this threshold down to 100 copies/mL [8]. The Roche **cobas**^®^HBV RNA investigational assay, developed for use on the high-throughput automated **cobas**^®^ 6800/8800 platforms, displays highly sensitive and reproducible measurement of cirB-RNA with a LOD less than 5 copies/mL and linear range of 10 to at least 10^7^ copies/mL in a broad range of HBV genotypes. This high sensitivity and genotype inclusivity is essential for monitoring antiviral drug efficacy and identification of patients at risk of reactivation when discontinuing antiviral treatment [19]. The fully automated cobas HBV RNA workflow enables high-throughput testing with minimal hands-on time. This assay does not require dilution of high viral loads samples to retain accuracy, contrary to the situation for ddPCR-based assays [8, 20].

Because of the common observation of high concentrations of HBV DNA in clinical samples, it is important to understand the degree to which RNA quantification by any molecular assay is impacted by DNA that contains the same target sequence. Our data demonstrate that RNA concentration measurements by cobas HBV RNA is not affected by the presence of up to 10^6^ times more DNA than RNA. Even at DNA:RNA ratios of approximately 10^7^, only modest differences were observed, which are unlikely to have clinical significance. In clinical practice it is extremely rare to observe DNA:RNA ratios greater than 10^5^ [21, 22].

Another automated assay for quantification of cirB-RNA has been described [9, 23]. Here we report precision and reproducibility with SD less than 0.15 log_10_ and 5% CV throughout the dynamic range. Given the different amplification strategies used by these two assays, head-to-head comparisons using paired serum and liver samples and more detailed determination of the RNA species detected will be of importance to determine which assay best reflects cccDNA load and transcriptional activity.

Differences in sample preparation and assay conditions (e.g. for nucleic acid amplification and detection) likely contribute to variability in quantitative molecular test results, which can be up to 100-fold using various WHO international standards [24–26]. The availability of an international standard is mandatory to properly compare results obtained from different studies [6, 7, 25]. The establishment of such a standard would enable a better interpretation of results generated with tests targeting different parts of the HBV genome. It is important to characterize how consistent, intact and stable the RNA component of the WHO DNA international standard may be, if it is intended to be used for HBV RNA standardization.

We demonstrated the utility of cobas HBV RNA using a panel of clinical samples encompassing a wide range of genotypes. We observed higher HBV DNA than cirB-RNA concentrations in untreated patients, similar to other reports [9, 16, 27, 28]. While it has been previously reported that HBV genotypes might have an influence on cirB-RNA levels [29], we did not observe any difference in RNA quantification or linearity based on genotype in our limited number of samples tested. Our data confirm previous reports that there are diverse patterns of cirB-RNA levels during the natural history of HBV infection, with higher cirB-RNA in HBeAg(+) patients indicating a higher degree of cccDNA transcriptional activity [15, 22, 30, 31].

In conclusion, the cobas HBV RNA investigational assay meets requirements related to automation, precision, sensitivity, specificity, linear range, and genotype inclusivity. Further studies in large cohorts of chronic hepatitis B patients, including clinical trials of drugs with novel modes of action aimed at HBV cure [5], are warranted to validate cobas HBV RNA as a tool for assessment of the clinical relevance of the cirB-RNA biomarker.

## Acknowledgments

This work is supported by the French National Research Agency Investissements d’Avenir program (cirB-RNA project – ANR-17-RHUS-0003). Presented in part at *The Liver Meeting* 2020 (Poster 737). Manuscript preparation services were provided by Data First Consulting (Sebastopol, CA) and funded by Roche Molecular Diagnostics.

## Author contributions

Writing—original draft: CS. Writing—review and editing: CS, MLP, BT, FZ, ML, AH, MH. Conceptualization: CS, AA, FZ, ML, MH. Patient inclusion MS, JF, FZ, ML. Data acquisition: CS, AB, CC, FB, MLP. Data analysis CS, BT, FZ, AA, ML, MH, MLP. Funding acquisition: ML, FZ, MH. All authors approved the final version to be submitted.

## Notes

### Competing Interest Statement

AH, BS, LW, AB, and MH are employees and stockholders of Roche Molecular Diagnostics. FZ has performed consulting activities for Aligos Therapeutics, Antios Therapeutics, Arbutus Biopharma, Assembly Bio, Enanta Pharmaceuticals, Enochian BioSciences, Gilead Sciences, Roche Molecular Diagnostics, Viravaxx AG, and received research funding via INSERM from Beam therapeutics. CS, CC, FB, JF, ML, BT, MLP declare no conflicts of interest.

## References

1. Fan R, Peng J, Xie Q, Tan D, Xu M, Niu J, et al., Combining Hepatitis B Virus RNA and Hepatitis B Core-Related Antigen: Guidance for Safely Stopping Nucleos(t)ide Analogues in Hepatitis B e Antigen-Positive Patients With Chronic Hepatitis B, J Infect Dis 222 (2020) 611–18. https://doi.org/10.1093/infdis/jiaa136

2. Fan R, Zhou B, Xu M, Tan D, Niu J, Wang H, et al., Association Between Negative Results From Tests for HBV DNA and RNA and Durability of Response After Discontinuation of Nucles(t)ide Analogue Therapy, Clin Gastroenterol Hepatol 18 (2020) 719–27.e7. https://doi.org/10.1016/j.cgh.2019.07.046

3. Farag MS, van Campenhout MJH, Pfefferkorn M, Fischer J, Deichsel D, Boonstra A, et al., Hepatitis B virus RNA as Early Predictor for Response to PEGylated Interferon Alfa in HBeAg Negative Chronic Hepatitis B, Clin Infect Dis (2020). https://doi.org/10.1093/cid/ciaa013

4. Ji X, Xia M, Zhou B, Liu S, Liao G, Cai S, et al., Serum Hepatitis B Virus RNA Levels Predict HBeAg Seroconversion and Virological Response in Chronic Hepatitis B Patients with High Viral Load Treated with Nucleos(t)ide Analog, Infect Drug Resist 13 (2020) 1881–88. https://doi.org/10.2147/IDR.S252994

5. Fanning GC, Zoulim F, Hou J, Bertoletti A, Therapeutic strategies for hepatitis B virus infection: towards a cure, Nat Rev Drug Discov 18 (2019) 827–44. https://doi.org/10.1038/s41573-019-0037-0

6. Charre C, Levrero M, Zoulim F, Scholtes C, Non-invasive biomarkers for chronic hepatitis B virus infection management, Antiviral Res 169 (2019) 104553. https://doi.org/10.1016/j.antiviral.2019.104553

7. Vachon A, Osiowy C, Novel Biomarkers of Hepatitis B Virus and Their Use in Chronic Hepatitis B Patient Management, Viruses 13 (2021). https://doi.org/10.3390/v13060951

8. Limothai U, Chuaypen N, Poovorawan K, Chotiyaputta W, Tanwandee T, Poovorawan Y, et al., Reverse transcriptase droplet digital PCR vs reverse transcriptase quantitative real-time PCR for serum HBV RNA quantification, J Med Virol (2020). https://doi.org/10.1002/jmv.25792

9. Butler EK, Gersch J, McNamara A, Luk KC, Holzmayer V, de Medina M, et al., Hepatitis B Virus Serum DNA andRNA Levels in Nucleos(t)ide Analog-Treated or Untreated Patients During Chronic and Acute Infection, Hepatology 68 (2018) 2106–17. https://doi.org/10.1002/hep.30082

10. Pasloske BL, Walkerpeach CR, Obermoeller RD, Winkler M, DuBois DB, Armored RNA technology for production of ribonuclease-resistant viral RNA controls and standards, J Clin Microbiol 36 (1998) 3590–4. https://doi.org/10.1128/JCM.36.12.3590-3594.1998

11. Stocher M, Berg J, Internal control DNA for PCR assays introduced into lambda phage particles exhibits nuclease resistance, Clin Chem 50 (2004) 2163–6. https://doi.org/10.1373/clinchem.2004.035519

12. ClinicalTrals.gov, Creation of a Cohort for the Quantitation and Characterization of Circulating Viral RNAs as a New Biomarker of Hepatitis B Functional Cure. (CirB-RNA). https://clinicaltrials.gov/ct2/show/NCT03825458. 2019 (accessed November 10).

13. European Association for the Study of the Liver, EASL 2017 Clinical Practice Guidelines on the management of hepatitis B virus infection, J Hepatol 67 (2017) 370–98. https://doi.org/10.1016/j.jhep.2017.03.021

14. Gao Y, Li Y, Meng Q, Zhang Z, Zhao P, Shang Q, et al., Serum Hepatitis B Virus DNA, RNA, and HBsAg: Which Correlated Better with Intrahepatic Covalently Closed Circular DNA before and after Nucleos(t)ide Analogue Treatment?, J Clin Microbiol 55 (2017) 2972–82. https://doi.org/10.1128/JCM.00760-17

15. Wang J, Yu Y, Li G, Shen C, Li J, Chen S, et al., Natural history of serum HBV-RNA in chronic HBV infection, J Viral Hepat 25 (2018) 1038–47. https://doi.org/10.1111/jvh.12908

16. Jansen L, Kootstra NA, van Dort KA, Takkenberg RB, Reesink HW, Zaaijer HL, Hepatitis B Virus Pregenomic RNA Is Present in Virions in Plasma and Is Associated With a Response to Pegylated Interferon Alfa-2a and Nucleos(t)ide Analogues, J Infect Dis 213 (2016) 224–32. https://doi.org/10.1093/infdis/jiv397

17. van Bommel F, Bartens A, Mysickova A, Hofmann J, Kruger DH, Berg T, et al., Serum hepatitis B virus RNA levels as an early predictor of hepatitis B envelope antigen seroconversion during treatment with polymerase inhibitors, Hepatology 61 (2015) 66–76. https://doi.org/10.1002/hep.27381

18. van Bommel F, van Bommel A, Krauel A, Wat C, Pavlovic V, Yang L, et al., Serum HBV RNA as a Predictor of Peginterferon Alfa-2a Response in Patients With HBeAg-Positive Chronic Hepatitis B, J Infect Dis 218 (2018) 1066–74. https://doi.org/10.1093/infdis/jiy270

19. Otsuka M, Koike K, Should Level of HBV RNA be Used to Determine When Patients Should Stop Treatment With Nucleos(t)ide Analogues, Clin Gastroenterol Hepatol 18 (2020) 551–52. https://doi.org/10.1016/j.cgh.2019.08.044

20. Nicot F, Cazabat M, Lhomme S, Marion O, Saune K, Chiabrando J, et al., Quantification of HEV RNA by Droplet Digital PCR, Viruses 8 (2016). https://doi.org/10.3390/v8080233

21. Ghany MG, King WC, Lisker-Melman M, Lok ASF, Terrault N, Janssen HLA, et al., Comparison of HBV RNA and Hepatitis B Core Related Antigen With Conventional HBV Markers Among Untreated Adults With Chronic Hepatitis B in North America, Hepatology 74 (2021) 2395–409. https://doi.org/10.1002/hep.32018

22. Zoulim F, Testoni B, Newsom C-L, Plissonnier M-L, Loglio A, Scholtes C, et al. Cross-sectional study of serum HBV RNA and HBcrAg in a real-life prospective cohort of 1500 chronic hepatitis B patients followed in France and Italy. The Liver Meeting (AASLD) 2021.

23. Anderson M, Gersch J, Luk KC, Dawson G, Carey I, Agarwal K, et al., Circulating Pregenomic Hepatitis B Virus RNA Is Primarily Full-length in Chronic Hepatitis B Patients Undergoing Nucleos(t)ide Analogue Therapy, Clin Infect Dis 72 (2021) 2029–31. https://doi.org/10.1093/cid/ciaa1015

24. Fryer JF, Minhas R, Dougall T, Rigsby P, Clare L, Morris CL, Collaborative Study to Evaluate the Proposed WHO 4th International Standard for Hepatitis B Virus (HBV) DNA for Nucleic Acid Amplification Technique (NAT) based assays. https://apps.who.int/iris/handle/10665/253052. 2016 (accessed December 7).

25. Baylis SA, Wallace P, McCulloch E, Niesters HGM, Nubling CM, Standardization of Nucleic Acid Tests: the Approach of the World Health Organization, J Clin Microbiol 57 (2019). https://doi.org/10.1128/JCM.01056-18

26. Baylis SA, Hanschmann KO, Schnierle BS, Trosemeier JH, Blumel J, Zika Virus Collaborative Study G, Harmonization of nucleic acid testing for Zika virus: development of the 1(st) World Health Organization International Standard, Transfusion 57 (2017) 748–61. https://doi.org/10.1111/trf.14026

27. Prakash K, Rydell GE, Larsson SB, Andersson M, Norkrans G, Norder H, et al., High serum levels of pregenomic RNA reflect frequently failing reverse transcription in hepatitis B virus particles, Virol J 15 (2018) 86. https://doi.org/10.1186/s12985-018-0994-7

28. Wang J, Shen T, Huang X, Kumar GR, Chen X, Zeng Z, et al., Serum hepatitis B virus RNA is encapsidated pregenome RNA that may be associated with persistence of viral infection and rebound, J Hepatol 65 (2016) 700–10. https://doi.org/10.1016/j.jhep.2016.05.029

29. van Campenhout MJH, van Bommel F, Pfefferkorn M, Fischer J, Deichsel D, Boonstra A, et al., Host and viral factors associated with serum hepatitis B virus RNA levels among patients in need for treatment, Hepatology 68 (2018) 839–47. https://doi.org/10.1002/hep.29872

30. Liu Y, Jiang M, Xue J, Yan H, Liang X, Serum HBV RNA quantification: useful for monitoring natural history of chronic hepatitis B infection, BMC Gastroenterol 19 (2019) 53. https://doi.org/10.1186/s12876-019-0966-4

31. Testoni B, Scholtès C, Plissonnier M-L, Berby F, Facchetti F, Villeret F, et al. Circulating HBV RNA correlates with intrahepatic covalently closed circular DNA (cccDNA) levels and activity in untreated chronic hepatitis B (CHB) patients. The International Liver Congress (EASL), (2021).

