## Supplemental Material for "Performance of the cobas^®^ HBV RNA Automated Investigational Assay for the Detection and Quantification of Circulating HBV RNA in Chronic HBV Patients"

### Supplementary Methods

#### Patient samples

Clinical samples were from 56 patients included in the ANR-17-RHUS-0003 cirB-RNA cohort (<https://clinicaltrials.gov/ct2/show/NCT03825458>), which is managed in accordance with the principles of the Declaration of Helsinki and the European guidelines for Good Clinical Practice and was approved by the independent Ethics Committees and registered with the French National Agency for Medicines and Health Products Safety (registration number: 2018-A02558-47). All samples were anonymized prior to the study, with an identification number containing no patient identifiers assigned to each sample. Upon reception, blood samples were centrifuged and aliquots of plasma or serum were stored at -80°C. Each aliquot was thawed and centrifuged before processing.

#### Analytical Specificity

Specificity was assessed using 20 HBV-negative serum and plasma samples and 20 replicates of a commercially available negative plasma sample (Basematrix, SeraCare, Milford, MA, USA). Cross-reactivity with other blood-borne viruses was assessed using remnant samples routinely tested at Lyon University Hospital containing human immunodeficiency virus type 1 (n=13, viral loads up to 6.1 log_10_ copies/mL), hepatitis C virus (n=10, viral loads up to 6.8 log_10_ IU/mL), or hepatitis E virus (n=5, with viral loads up to 6.3 log_10_ copies/mL).

#### Matrix equivalency

Ten paired serum and plasma samples, each from the same patient were tested. HBV DNA concentrations ranged from 2 to 6 log_10_ IU/mL).

#### Digital droplet PCR (ddPCR) assays

arRNA was quantified using the One-step RT-ddPCR Advanced kit: 22-μL reaction mixture was prepared comprising 5 μL of Supermix™ for probes, 2 μL of reverse transcriptase enzyme (20 U/μL), 1.1 μL of in-house designed primer and probe mix located at the 3’ end of HBV transcripts (nt 1780-1848 referred to EcoRI site; strain ayw, U95551.1: Fwd-GGCTGTAGGCATAAATTGGTC, Rev-CAAGAGATGATTA GGCAGAGG, [6FAM]CGCACCAGCACCATGCAACTTT[BHQ1]), 1 μL of DTT (15 mM) and 5 μL of arRNA. Droplet formation was carried out using a QX200 Automated droplet generator. Reverse transcription step was performed at 50°C followed by 95 °C inactivation for 10 min.

PCR amplification was performed in the C1000 Touch™ Thermal cycler (Bio-Rad) with a ramp rate of 2 °C/s and the lid heated to 105 °C, according to Bio-Rad recommendations. PCR comprised 40 cycles of denaturation at 95 °C for 30 s and 60 °C for 1 min. Final inactivation was performed for 10 minutes at 98°C and the reaction was kept at 4°C. Subsequently, a droplet reader was used to calculate the number of both positive and negative droplet events from each PCR reaction mixture. The number of RNA copies were determined by calculating the ratio of the positive droplets over the total droplets combined with Poisson distribution by QuantaSoft analysis software.

#### HBV RNA quantitative RT-PCR assay (manual version)

Total RNA was isolated from 200 µL of plasma using a cell free RNA sample preparation method. Briefly, samples were diluted in 300 µL of elution buffer prior to incubation in lysis buffer supplemented with proteinase K, included in the kit, for 30 minutes at room temperature (15 - 30°C). After addition of ethanol to bind the RNA, the lysate was transferred to a High Pure Extender Assembly unit (Roche Diagnostics) and centrifuged at 2000 g for 5 minutes. A DNase treatment was performed at room temperature for 15 minutes. Extensive washes were performed and RNA was eluted in 40 µL of RNA elution buffer.

HBV RNA amplification and quantification were performed using a manual workflow version of the HBV RNA assay. Briefly, the working mastermix was prepared by adding 10 µL per reaction manganese acetate solution (16.5 mM) to the HBV RNA assay mastermix. One microliter of a Generic Internal Control armored RNA was added for each sample. For the reaction, 20 µL of sample RNA (input) was added to 30 µL of working mastermix with GIC. Amplification must be started within 1 hour from the time that the processed samples and controls are added to the working mastermix. Quantitative PCR was performed on the Light Cycler® 480 II analyzer (LC480II, Roche Diagnostics) for the target (HEX dye: 533 excitation-580 emission) and the GIC (CY5.5 dye: 618 excitation-660 emission. Copy number was calculated based on standard curve generated with arRNA.

#### Statistical Analysis

The relationship between RNA concentrations measured in different assays was assessed by Deming regression and Bland–Altmann analysis (Graphpad Prism version 9).

### Supplementary results

##### Table S1. Patient sample characteristics

|  |  | Untreated ^a^ | NUC-treated | Total |
| --- | --- | --- | --- | --- |
|  | N | 36 | 20 | 56 |
|  | Age range (years) | 19-77 | 19-72 | 19-77 |
| Genotype | A | 3 | 2 | 5 |
|  | B | 4 | 2 | 6 |
|  | C | 4 | 3 | 7 |
|  | D | 11 | 4 | 15 |
|  | E | 12 | 4 | 16 |
|  | F | 1 | 2 | 3 |
|  | G | 0 | 1 | 1 |
|  | unknown | 1 | 2 | 3 |
| HBeAg status | positive | 13 | 8 | 21 |
|  | negative | 23 | 12 | 35 |
| HBV DNA | >LLOQ | 36 | 12 | 48 |
|  | <LLOQ, >LOD | 0 | 5 | 5 |
|  | <LOD | 0 | 3 | 3 |
|  | not done | 0 | 1 | 1 |
|  | mean HBV DNA (log_10_ IU/mL) ^b^ | 5.49 | 2.97 | 4.86 |
| HBV RNA | >LLOQ | 25 | 9 | 34 |
|  | <LLOQ, >LOD | 8 | 3 | 11 |
|  | <LOD | 3 | 8 | 11 |
|  | not done | 0 | 0 | 0 |
|  | mean HBV RNA (log_10_ copies/mL) ^b^ | 4.21 | 3.37 | 3.99 |

^a^ including one with unknown treatment status

^b^ mean of results > LLOQ

##### Table S2. Patient samples used for linearity testing

| Genotype | Disease phase^a^ | HBeAg  status | Treatment | HBV DNA  (log_10_ IU/mL) | cirB-RNA  (log_10_ copies/mL) |
| --- | --- | --- | --- | --- | --- |
| A | CH | + | No | 8.8 | 6.1 |
| B | CI | + | No | 8.3 | 6.5 |
| C (1) | CH | + | Yes | 1.7 | 6.8 |
| C (2) | CH | + | No | 8.3 | 6.2 |
| D (1) | CH | + | Yes | 4.5 | 6.3 |
| D (2) | CI | + | No | 8.2 | 5.9 |
| E | CI | + | No | 8.7 | 5.9 |

^a^ CH: chronic hepatitis; CI: chronic infection

##### Table S3. Sensitivity confirmation using clinical samples

| **Genotype** | **Nominal concentration (copies/mL)** | **N** | **N detected** | **% detected** |
| --- | --- | --- | --- | --- |
| A | 14.7 | 2 | 2 | 100% |
|  | 7.4 | 5 | 4 | 80% |
|  | 3.7 | 5 | 5 | 100% |
|  | 1.8 | 5 | 2 | 40% |
| B | 15.6 | 1 | 1 | 100% |
|  | 10.0 | 1 | 1 | 100% |
|  | 7.8 | 2 | 2 | 100% |
|  | 3.9 | 5 | 3 | 60% |
| C | 36.7 | 2 | 2 | 100% |
|  | 18.4 | 5 | 5 | 100% |
|  | 9.2 | 5 | 5 | 100% |
|  | 4.6 | 5 | 5 | 100% |
| D | 15.8 | 2 | 2 | 100% |
|  | 7.9 | 5 | 5 | 100% |
|  | 4.0 | 5 | 4 | 80% |
|  | 2.0 | 5 | 2 | 40% |
| E | 25.0 | 2 | 2 | 100% |
|  | 12.5 | 5 | 5 | 100% |
|  | 6.3 | 5 | 5 | 100% |
|  | 3.1 | 5 | 4 | 80% |

#### Precision and reproducibility.

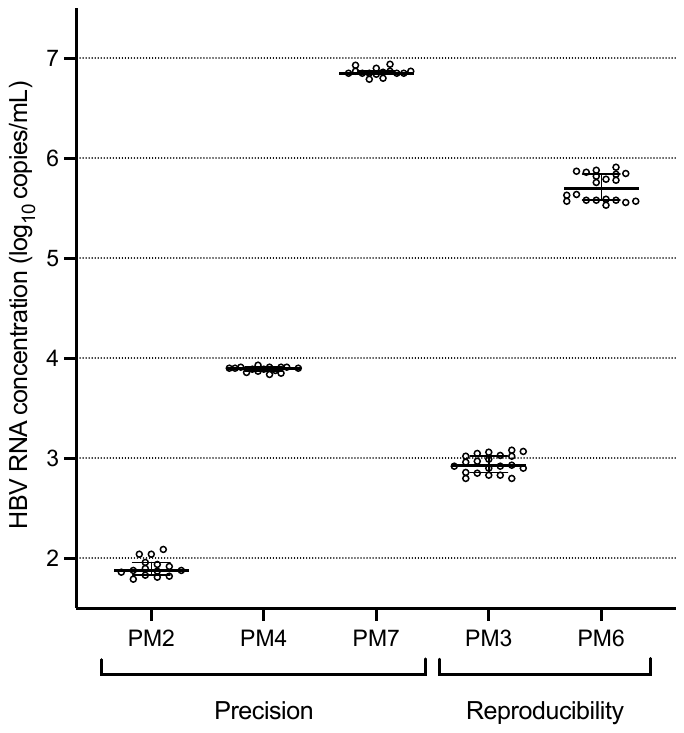

##### Figure S1. Precision and Reproducibility results. Replicate results for each panel member (PM) are shown. Horizontal bars represent the median and error bars represent the 95% confidence intervals.

#### Matrix equivalency

The mean difference in RNA titer between paired serum and plasma samples was at 0.22 ± 0.27 log_10_ copies/mL (Figure S2). Both specimen types are thus confirmed to be equivalent and suitable for this assay.

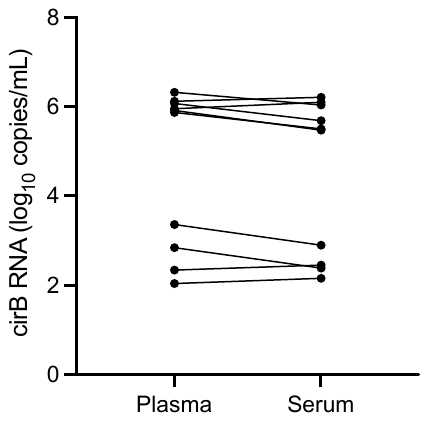

##### Figure S2. Quantification of cirB-RNA in plasma and serum. Lines connecting the data points link samples from the same patient.

#### Method Comparison: cobas HBV RNA vs. an in-house ddPCR assay

There was excellent concordance between arRNA concentrations measured with cobas HBV RNA and the in-house ddPCR assay (Figure S3).

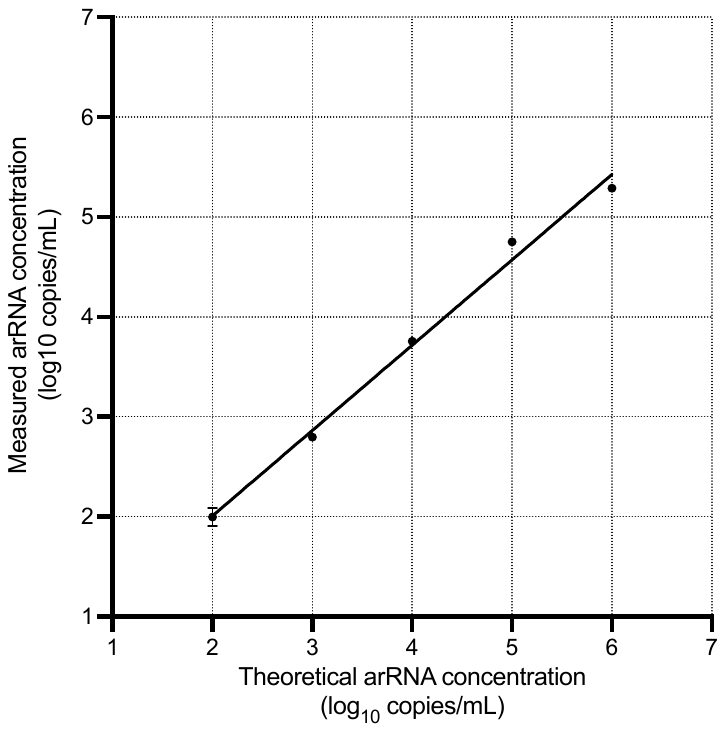

##### Figure S3. Method comparison (cobas HBV RNA vs. in-house ddPCR assay) with arRNA. The average of duplicate results are plotted.

#### Method Comparison: cobas HBV RNA vs. manual workflow

For research purposes that do not require high throughput and automation, a manual version of the HBV RNA assay was developed. Results obtained with this manual assay were highly comparable to those obtained with cobas HBV RNA (Figure S4).

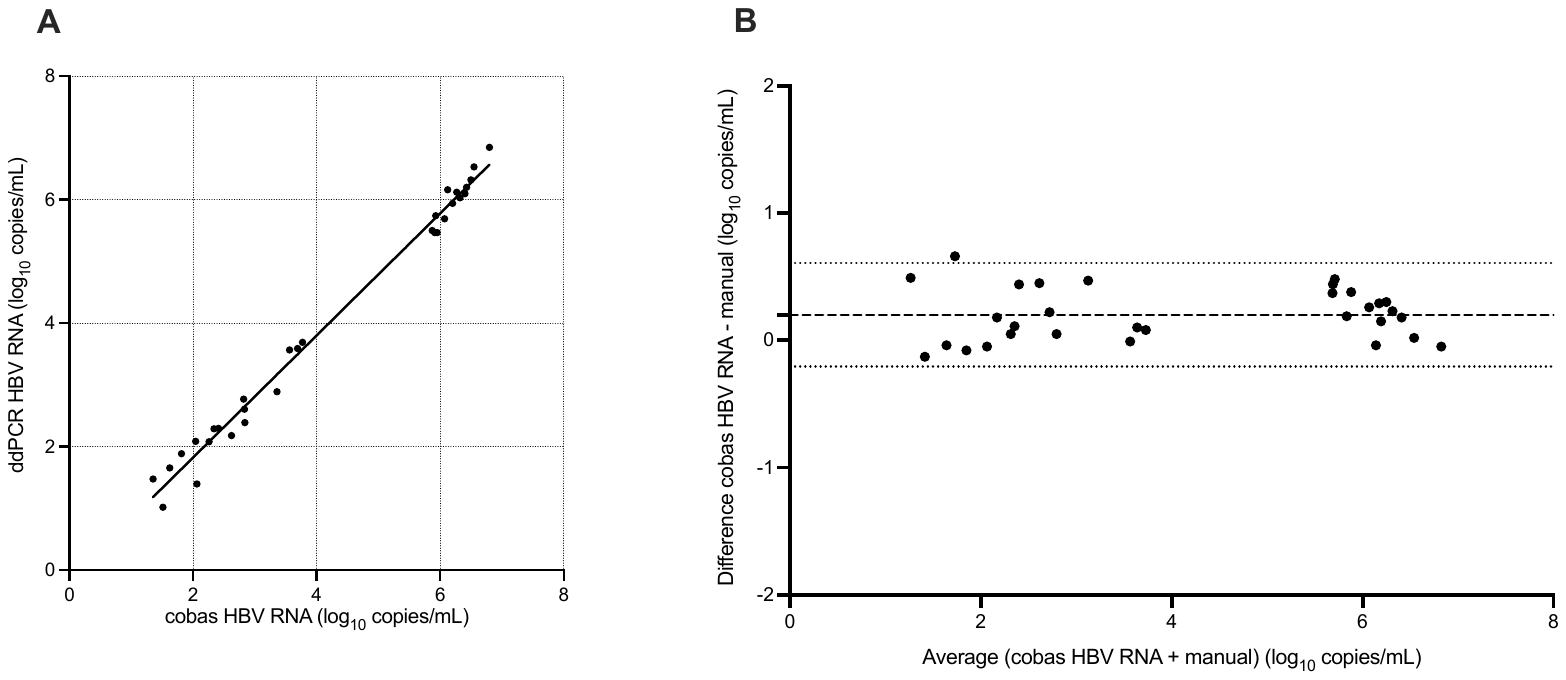

##### Figure S4: Comparison between cobas HBV RNA and manual assay. (A) Demming regression. (B) Bland-Altman plot. The average difference and limit of agreement are shown by dashed line and dotted lines, respectively.

#### Evaluation of patient samples

We measured cirB-RNA levels in 56 clinical samples, from 36 untreated and 20 NUC-treated, HBV-infected patients. The samples were selected for their genotype representativity (A to G) and to cover a wide HBV DNA viral load range within the different phases of HBV disease according to EASL guidelines [1] (Figure S5).

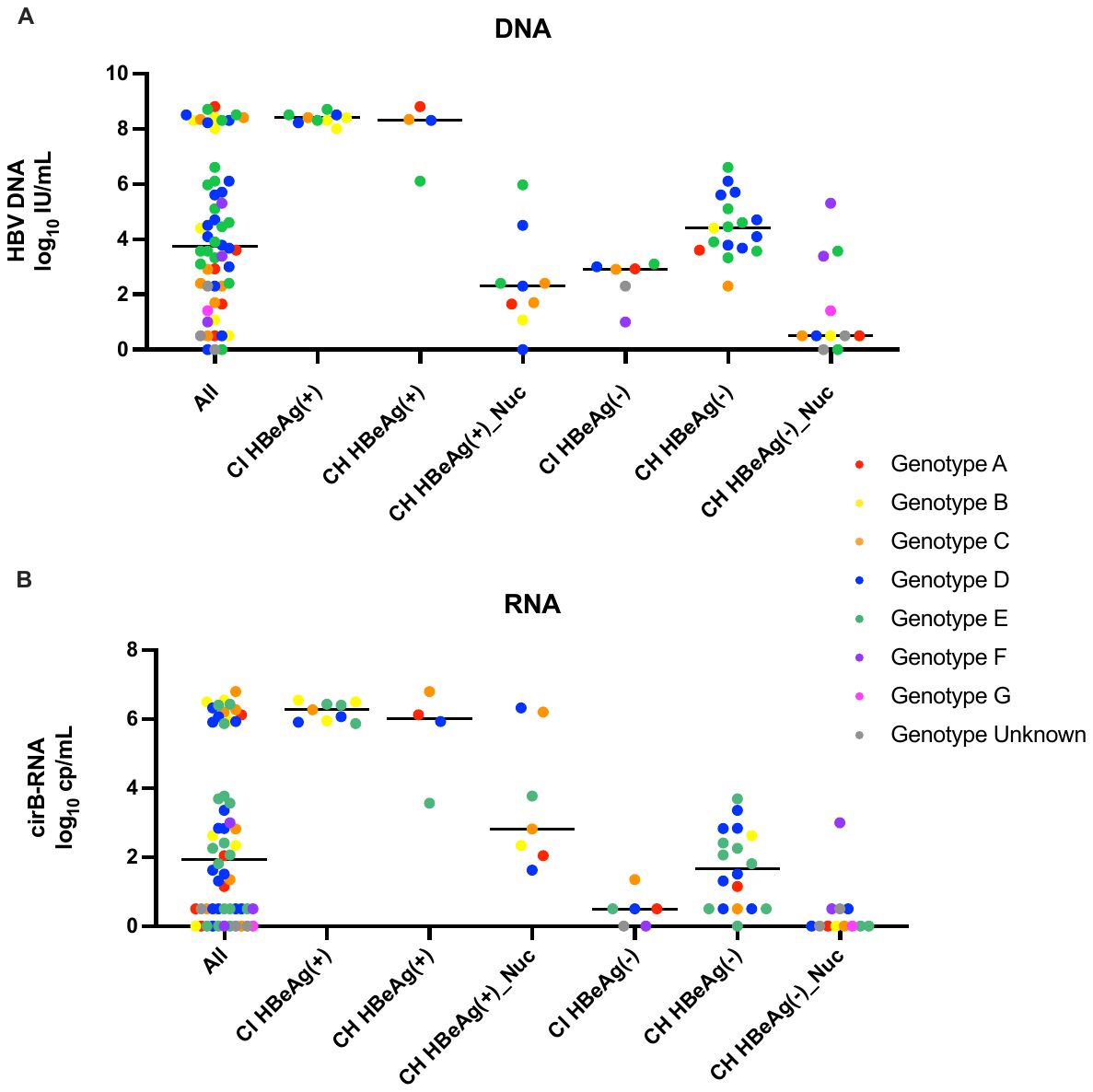

##### Figure S5. Comparison of HBV DNA and RNA concentrations in patient samples

### References

1. European Association for the Study of the Liver, EASL 2017 Clinical Practice Guidelines on the management of hepatitis B virus infection, J Hepatol 67 (2017) 370-98. <https://doi.org/10.1016/j.jhep.2017.03.021>
